# Viral haplotype reconstruction from long reads with virCHap

**DOI:** 10.64898/2026.02.09.704753

**Authors:** Yun Gao, Ting Yu, Bingqiang Liu, Guojun Li

## Abstract

Resolving genomes at the haplotype level for viral populations is crucial for understanding the prevalence of viral diseases and for the development of effective therapeutic treatments. However, viral haplotype reconstruction still presents challenges, such as an unknown number of strains, high inter-strain similarity, repetitive regions, and difficulties in abundance estimation. Here, we developed virCHap, a new reference-based haplotype phasing algorithm for viruses, which applies graph partitioning followed by iteratively quantifiable cluster merging on long-read sequencing data. Benchmarking on simulated and real datasets demonstrates that virCHap outperforms current tools in terms of recall, accurate abundance estimates and read clustering accuracy. On the simulated large-genome VZV experiment, virCHap had a 96.5% recall, 14% higher than the second-best method, and had the most accurate abundance estimates. On a real 5-strain PVY dataset, virCHap had a precision exceeding 92.9%, a recall of over 97%, and a read clustering accuracy of 82%, outperforming the second-best method by 32%. On a real 6-strain SARS-CoV-2 dataset, virCHap achieved >96.9% accuracy, and the most accurate abundance estimates within the spike gene.

## Introduction

Many viruses, including RNA viruses such as influenza, HIV, hepatitis C, and SARS-CoV-2, as well as certain DNA viruses like hepatitis B, possess relatively high mutation rates due to their error-prone replication process and the lack of sophisticated repair mechanisms [1]. The high mutation rates result in viral populations that exist as a collection of closely related strains that differ by only small amounts of variants, which are called quasispecies [2]. The presence of quasispecies enables viruses to readily adapt to dynamic environments and develop resistance to antiviral drugs and vaccines, making the design of effective, long-lasting treatments for viral diseases exceedingly difficult [3]. Determining the strain sequences and estimating their prevalence using sequencing reads for viral populations help the understanding of viral diseases and provides guidance for developing effective medical therapeutics.

Resolving virus genomes at the strain haplotype level, i.e., a sequence of variants observed jointly and in sufficient abundance by sequencing reads, provides critical insights into quantifying the genetic diversity of viral quasispecies and elucidating viral transmission, evolution, and fitness [4, 5]. For instance, during the early stages of the COVID-19 pandemic, genomic analysis of cases in Guangdong Province, China, successfully distinguished imported cases from subsequent local transmission chains, clearly revealing the early transmission dynamics [6]. Despite the short length of viral genomes, viral haplotype reconstruction remains challenging due to the unknown number of strains, high sequence similarity among strains, the presence of repetitive genomic regions, and difficulties in abundance estimation.

The development of sequencing technologies has significantly advanced the progress of strain-level viral reconstruction. Haplotype reconstruction methods can be broadly categorized into two types: de novo haplotype assembly and reference-based haplotype phasing. De novo assemblers typically are used to generate a species-level consensus, suppressing strain heterogeneities, as assembly algorithms, such as Canu [7], Flye [8], or Shasta [9], were originally designed for error-prone long reads with error rates above 10%. Recently developed PacBio HiFi assembly algorithms, such as hifiasm [10], HiCanu [11], hifiasm-meta [12], and metaMDBG [13], take advantage of highly accurate reads to use (rather than suppress) heterozygous variants to generate haplotype assemblies, however, HiFi data are not always available, and the assembly process itself is computationally intensive [14]. Haplotype phasing can be used to complement de novo assembly, and leveraging information from the known virus often leads to more complete haplotypes [15]. Reference-based haplotype phasing identifies co-occurring variants of different strains within the same species over variant sites from sequencing data aligned to the reference.

We focus on viral haplotype phasing by using third-generation sequencing data. Although next-generation sequencing (NGS) reads (e.g., Illumina reads) are more accurate, NGS reads are short, which is generally under 300 bp [16], making full-length haplotype reconstruction difficult. Moreover, they are often too short to span multiple heterozygous sites consistently across longer distances, and thus they cannot well distinguish between inter-genomic and intra-genomic highly similar regions. In contrast, long sequencing reads can overcome these limitations. The long sequencing reads can span more variant positions, directly enabling the long phasing of variants within individual reads, and improving the resolution of repetitive regions. However, the high error rate of raw long reads introduces new challenges for accurate haplotype reconstruction.

The earliest publicly available software tool for viral haplotype phasing is ShoRAH [17], which is used for the analysis of NGS reads by integrating a path cover-based approach with probabilistic clustering [18]. aBayesQR [18] uses a maximum-likelihood framework to infer individual sequences within a mixture from NGS data. CliqueSNV [19] is designed for both NGS and PacBio data, leveraging linkage between SNVs to construct an SNV graph and then merging cliques in the graph to generate haplotypes. iGDA [20], RVHaplo [21], HaploDMF [22], and devider [14] are haplotype phasing algorithms specifically developed for long-read data. iGDA detects minor SNVs using maximum conditional probability, followed by the Adaptive-Nearest Neighbor (ANN) clustering algorithm to cluster reads and obtain haplotypes. RVHaplo identifies SNVs through two binomial tests and maximum conditional probability, then clusters the reads to reconstruct haplotypes. HaploDMF applies a deep matrix factorization model along with a clustering algorithm to group reads and generate haplotypes. devider resolves haplotypes by finding paths in a positional de Bruijn graph (PDBG). Strainline [23] is a de novo method that takes the longest reads as templates, aligns other reads against them to produce error-corrected contigs, and subsequently constructs haplotypes based on overlaps among the contigs.

Here, we present virCHap, a new reference-based algorithm for haplotype phasing of **vir**us genomes from long-read sequencing data. It consists of two stages: **c**lustering and merging clusters, to obtain strain **hap**lotypes, where the first clustering using the label propagation algorithm [24] is based on the similarity between reads and then the results are merged to obtain longer haplotypes. virCHap does not require prior knowledge of the number of strains. We validated the performance of virCHap on three simulated and three real datasets. The evaluation results demonstrated that virCHap showed significant improvement over the other methods, including devider, HaploDMF, iGDA, RVHaplo and Strainline, in terms of recall, accurate abundance estimates and read clustering accuracy.

### Overview of the virCHap

virCHap requires a reference genome, a set of aligned reads and SNPs as input and then outputs the phased SNPs, the clustered reads, the base-level sequences and the abundance estimate for each strain haplotype (**Fig. 1**). The long reads aligned to the reference are represented as a set of heterozygous SNPs, hereafter referred to as reduced reads. First, virCHap builds an overlap graph, which encodes the pairwise distances between the reduced reads. It then performs clustering in two main stages: initial partitioning followed by iterative merging. virCHap uses the label propagation algorithm to partition the graph into densely connected clusters. A consensus sequence is computed for each cluster, after which they are evaluated for merging. Two clusters are merged only if the newly formed one shows minimal deviation from its originals; otherwise, merging is not performed. This merging process repeats iteratively until no further merges are possible.

**Fig. 1.**
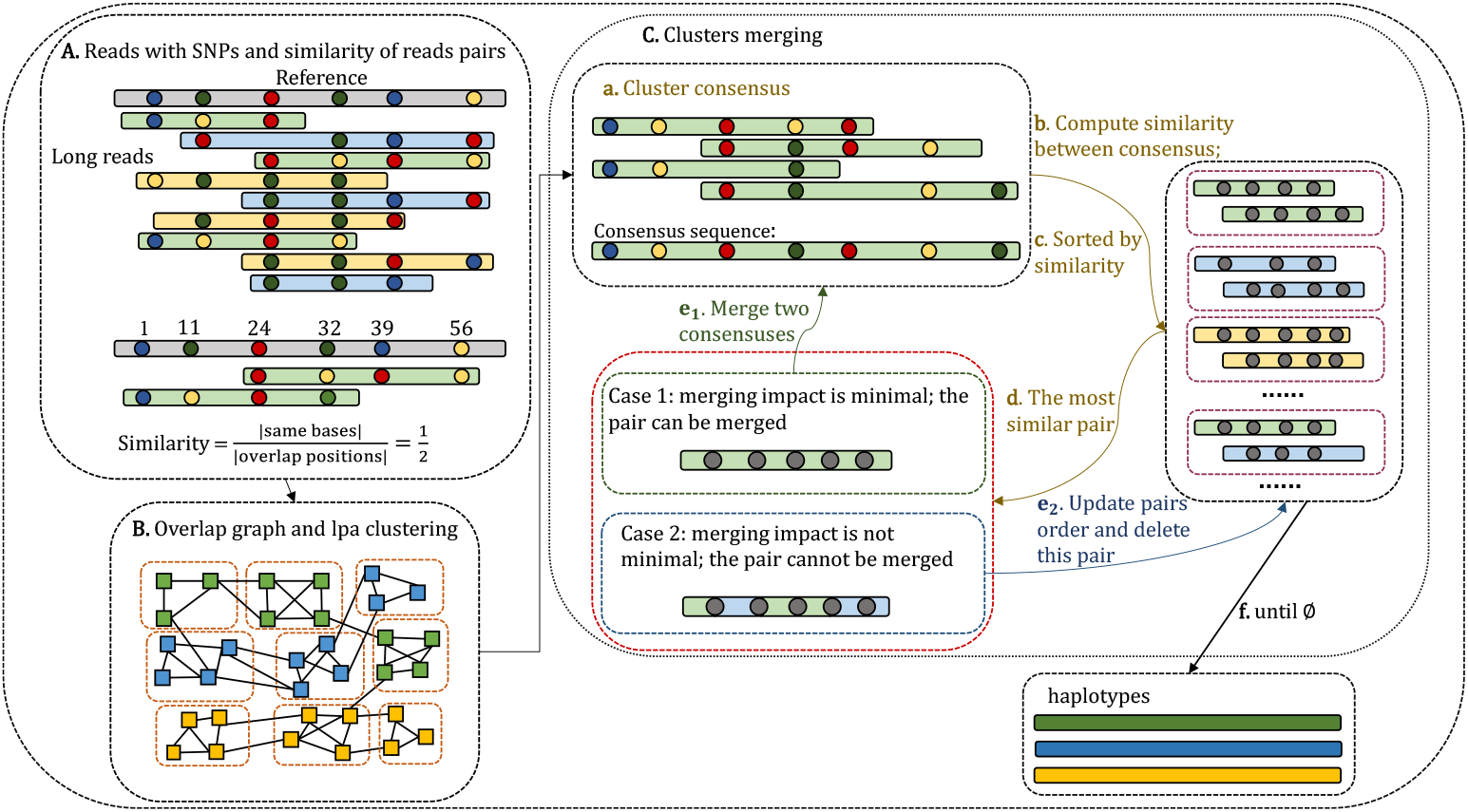
Overview of the virCHap workflow. **A**. Reads that are aligned to a reference are converted to a set of SNPs named as reduced reads. The colored dots are SNP bases. The similarity between reduced reads is defined as the number of the same SNPs divided by the number of overlapped SNP positions. **B**. The overlap graph reflecting pairwise read similarity is built by reduced reads and is partitioned by the label propagation algorithm. **C**. The iterative process of merging clusters. (**a**) Convert clusters to consensus sequences. (**b**) Calculate pairwise similarity between consensus sequences; (**c**) Sort the similarities in descending order; (**d**) Select the most similar pair. If the two clusters can be merged (**e**_**1**_), compute the consensus sequence for the merged cluster and update the similarity set; if not (**e**_**2**_), remove this pair and update the similarity set. The iteration continues until the similarity set is empty, and the final haplotypes are generated (**f**).

## Results

Because different viruses have different replication rates and intrasample similarities [22], we assessed virCHap on various viruses, including hepatitis C virus (HCV), human immunodeficiency virus (HIV), varicella-zoster virus (VZV), norovirus, potato virus Y (PVY), and severe acute respiratory syndrome coronavirus 2 (SARS-CoV-2). We evaluated virCHap on both simulated and real datasets. In the simulated experiments, we first tested virCHap on the datasets with varying coverages from 9 HCV strains and then on the datasets with different numbers of strains from 20 HIV strains. Next, we tested on longer genomes using 6 VZV strains (∼125 kb), including datasets with both uniform and non-uniform abundances. For all simulated datasets, the long reads were simulated using Badread [25]. The long reads were subsequently mapped to the reference by minimap2 [26], and variants were called using LoFreq [27]. In the real data experiments, we evaluated virCHap on seven norovirus strains dataset, five PVY strains dataset, and six SARS-CoV-2 strains dataset. The long reads were aligned to the reference by minimap2 and variants were called by LoFreq. Detailed commands are provided in the “Command lines for replicating the analyses” of the Supplementary Materials.

We benchmarked virCHap v1.0 against devider v0.0.1, HaploDMF (v May 2022), iGDA v1.01, RVHaplo v2 and Strainline (v Mar 2021). We ran all methods for their default settings with 4 threads. We compared the predicted haplotypes against the truth haplotypes and got the best-matchings between predicted haplotypes and truth haplotypes for evaluation. The following metrics were used for benchmarking:

**precision**: precision = TP / (TP + FP), where TP = true positive; the number of SNPs is attributed to the correct haplotypes. FP = false positive; the number of SNPs does not belong to this haplotype.

**recall**: recall = TP / (TP + FN), where FN = false negative; the number of SNPs is not represented in the results.

**read clustering accuracy**: the percentage of reads that were correctly attributed to their truth haplotypes.

**NGA50**: The length N that 50% of the reference sequence is covered in contigs with length ≥ N after breaking contigs at misassembly events and removing all unaligned bases. The reference sequence here is the concatenation of all strain-specific genomes. NGA50 is reported by MetaQUAST v5.3.0 [28] with the option --unique-mapping.

**Earth mover’s distance (EMD)** [14]: a measure of the distance between the predicted abundances and the true abundances.

**nContigs**: the number of predicted haplotypes.

### Evaluation on simulated HCV data

Hepatitis C virus (HCV) is grouped in the genus Hepacivirus within the family *Flaviviridae*, and it has a positive-sense RNA genome and high genomic variability [29]. We downloaded ten haplotypes (9273-9311bp) [30] of HCV, Subtype 1a, from the NCBI database, which were used in Strainline. We randomly chose a haplotype with accession EU255989 as the reference and the remaining nine haplotypes were used to simulate reads by Badread under a specified abundance ratio 1:3:7:9:12:15:20:25:30, with the smallest strain coverage ranging from 3x to 100x and an average read length of 6000 bp. The pairwise divergence ranges from 2.8% to 7.4% (**Supplementary Fig. 1a**), which is derived from 1 - ANI, where average nucleotide identity (ANI) is calculated using FastANI [31].

As shown in **Fig. 2**, the performance of all methods improved as the coverage increased, with the recall metric exhibiting a more rapid improvement. The mean precision of HaploDMF, devider, virCHap, and iGDA all exceeded 99%. virCHap’s mean recall (96.4%) was slightly lower (1.16%) than the top-performing HaploDMF, but 4.63% higher than the third-best devider. virCHap had the highest mean read clustering accuracy (98.8%) versus HaploDMF (96.9%; the second best). HaploDMF, virChap and devider consistently yielded higher NGA50, all larger than 9290bp, and produced the number of predicted haplotypes closer to the true number of strains (the 9), demonstrating strong contiguity. Additionally, these three methods achieve the smallest distance between predicted and true abundance profiles compared to the other methods. As shown in **Fig. 2**, HaploDMF and virCHap produced robust outputs even under low coverage conditions.

**Fig. 2.**
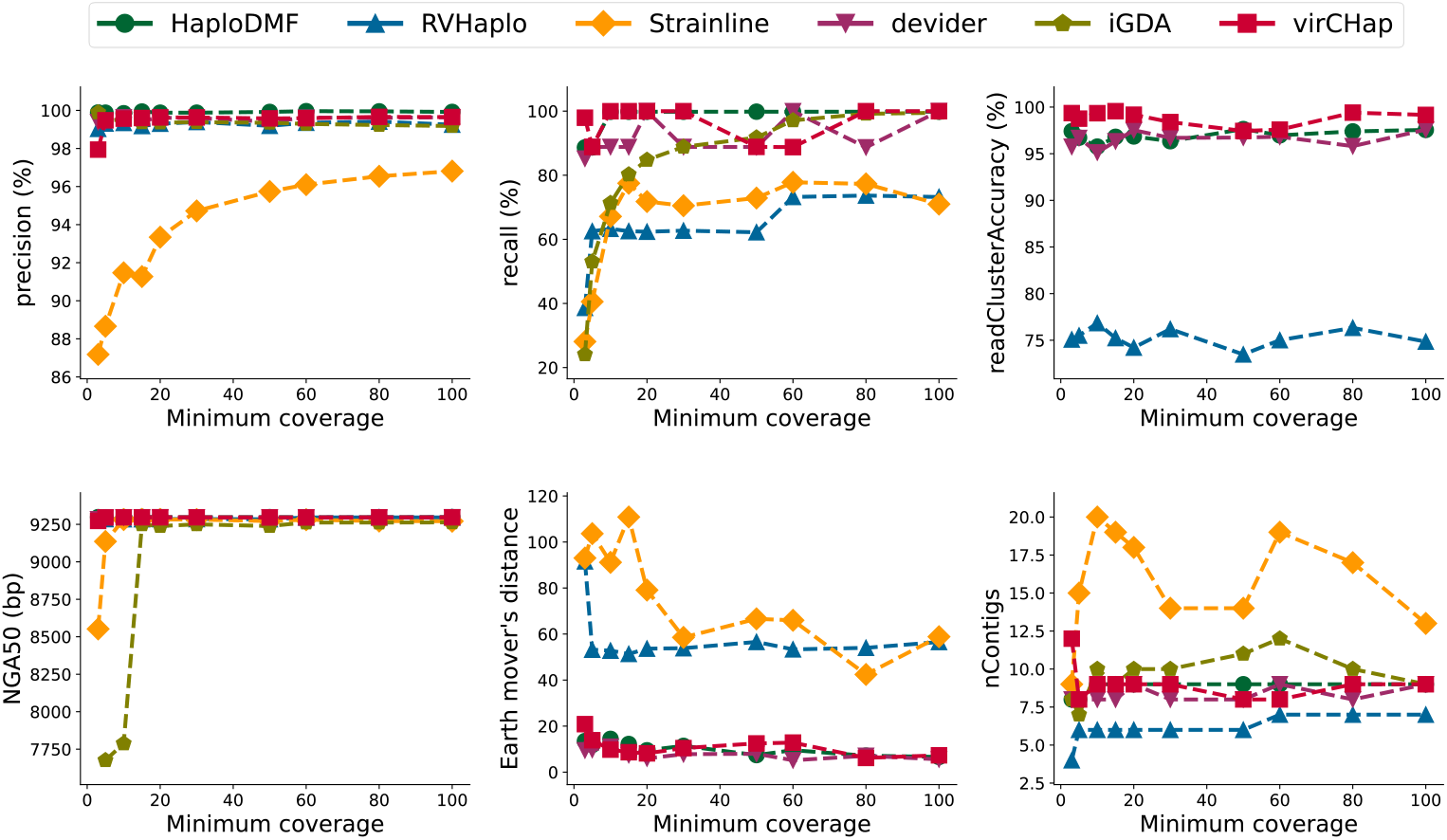
Benchmarking results for six methods on the simulated HCV datasets. 9 HCV strains with abundance ratios 1:3:7:9:12:15:20:25:30 and the lowest strain coverage ranging from 3x to 100x. The x-axis indicates the coverage for the lowest-coverage strain.

### Evaluation on simulated HIV data

To assess the performance of various methods across different numbers of strains, we selected 20 HIV haplotypes [32]. The pairwise divergence among these haplotypes ranges from 0.3% to 8.9% (**Supplementary Fig. 1b**). We simulated reads for 2 to 20 strains using Badread, with an average read length of 6000 bp. The abundance ratios increased proportionally to the number of strains (e.g., an abundance ratio of 1:2:3:4 for 4 strains) and the smallest strain coverage was set to 10x. The sequence with accession OR483991 (8971bp) was used as the reference.

iGDA did not output results on the dataset with 2 strains, therefore, mean values of each metric for iGDA were computed only across the datasets where outputs were available. As shown in **Fig.3**, as the number of strains increased, the performance of most methods generally declined. While precision remained stable with strain number increasing, recall, EMD, and read clustering accuracy showed marked deterioration, except for iGDA (**Fig. 3**). virCHap had the highest mean recall (71.7%), compared to devider (63.9%, the second highest) and HaploDMF (54.3%, the third highest). virCHap had slightly lower mean precision (97.9%) than iGDA (99.3%), HaploDMF (99.0%), and RVhaplo (98.7%), but was slightly higher than devider (97.4%). Moreover, virCHap had the highest mean read clustering accuracy (83.8%) versus devider (76.2%, the second best). Owing to the relatively low pairwise divergence among some of the 20 HIV strains (**Supplementary Fig. 1b**), accurate discrimination between haplotypes was challenging, leading to generally low recall across all methods. The number of predicted haplotypes for RVHaplo, devider, HaploDMF, and virCHap was nearly all lower than the truth number of strains. As the number of strains increased, the difference between the number of predicted haplotypes for iGDA and the true number of strains transitioned from negative to positive, accompanied by an increase in recall, however, its recall remained relatively low and it had the lowest mean NGA50. Strainline generated a high number of haplotypes with large mean NGA50, which indicated high contiguity, however, it had the lowest mean precision (90.9%), a lower recall (44.5%) and the worst mean EMD (68.9).

**Fig. 3.**
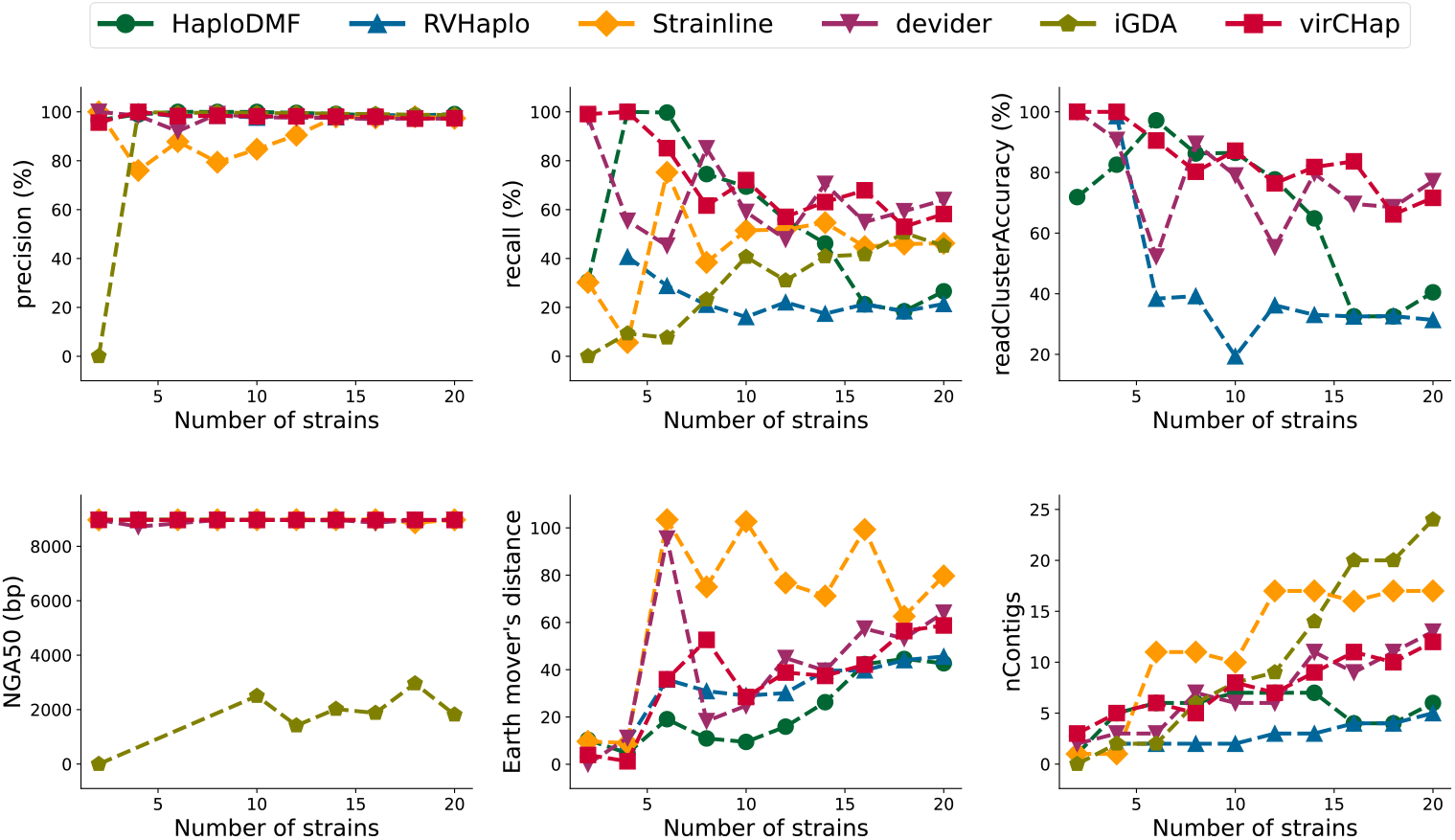
Benchmarking results for six methods on the simulated HIV datasets. 2 to 20 HIV strains with 30x – 210x coverage and arithmetically increasing abundance ratios (e.g., an abundance ratio of 1:2:3:4 for 4 strains). The x-axis indicates the number of strains.

### Evaluation on simulated varicella-zoster virus (VZV) data

Varicella-zoster virus (VZV), a member of the subfamily *Alphaherpesvirinae* within the *Herpesviridae* family, is a highly conserved double-stranded DNA virus that exclusively infects humans. VZV is one of the largest viruses with a genome size of ∼125 kb. The variation between different strains is manifested as single nucleotide polymorphisms (SNPs) [33]. To access the performance of various methods on large viral genomes, we selected six varicella-zoster virus (VZV) strain haplotypes from the NCBI database. The pairwise divergence among these haplotypes ranges from 0.05% to 0.25% (**Supplementary Fig. 1c**). We simulated two types of datasets for 2-6 strains: one with abundance ratios increasing proportionally to the number of strains (e.g., a ratio of 1:2:3:4 for 4 strains), where the smallest strain coverage was set to 10x; and another with uniform abundance, where each strain coverage was set to 30x. The sequence with accession GCF_000858285.1 (124884 bp) was used as the reference.

Since Strainline and iGDA produced no output on some datasets, the mean values of each metric for each method were computed only across the datasets where their results were available. As shown in **Fig. 4**, virCHap had the highest mean recall (96.5%), exceeding the second-best, HaploDMF, by 14.2%, though virCHap’s mean precision (96.6%) was 2.4% lower than the best, iGDA. virCHap had the highest read clustering accuracy (96.4%), exceeding the second-best method, RVHaplo, by 11.1%. Additionally, virCHap had the lowest mean EMD (11.0 vs. 15.8 of RVHaplo, the second best). Except for iGDA, which predicted more than three times the true number of haplotypes in uniform coverage datasets with 5 and 6 strains, all other methods generated haplotype numbers close to the truth strain number. The mean NGA50 for all methods was no less than 110 kb. Besides, as shown in **Fig. 4**, iGDA, devider, and virCHap generally performed better on uniform abundance datasets than on non-uniform ones, whereas HaploDMF had the opposite trend, performing better on non-uniform abundance datasets than on uniform ones.

**Fig. 4.**
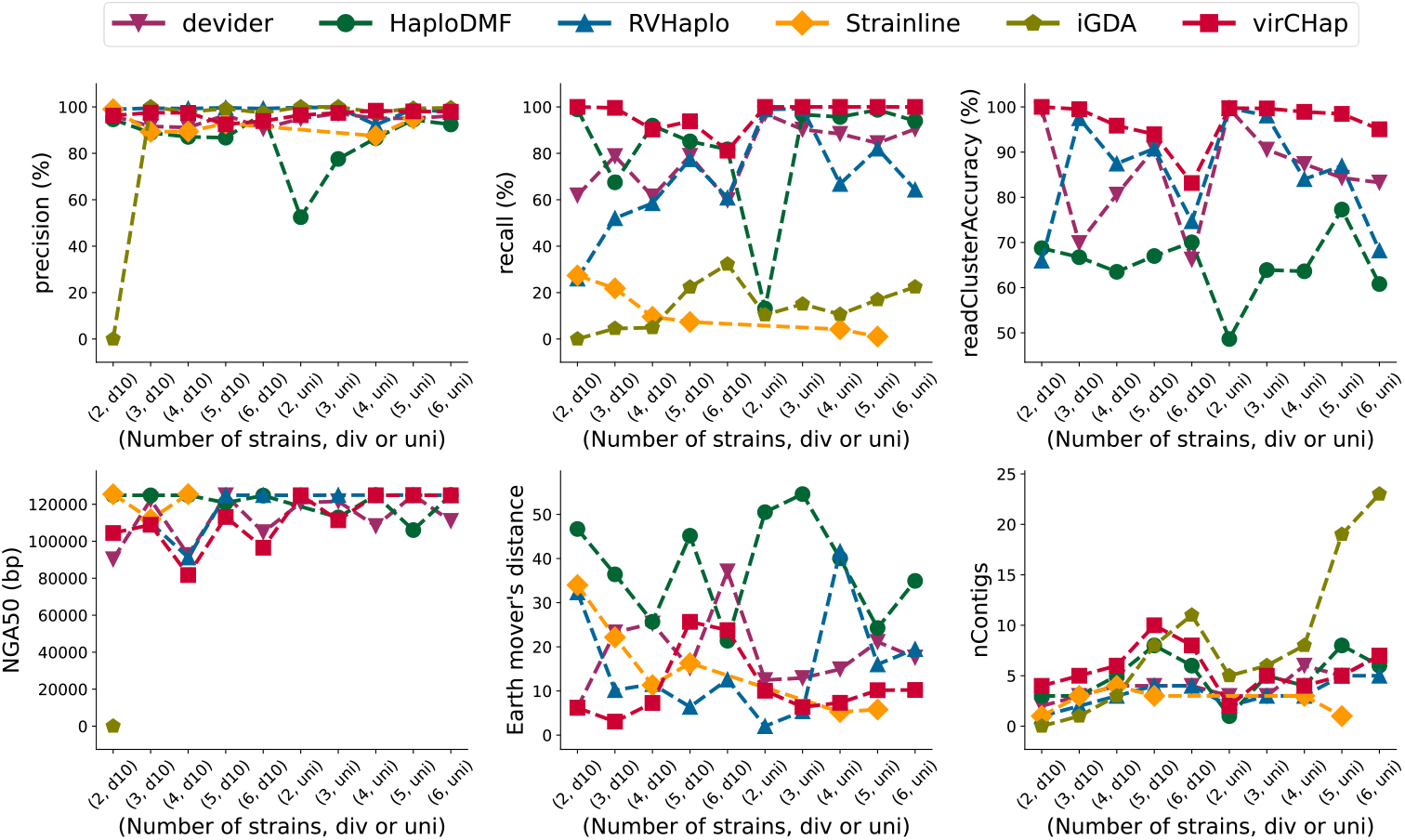
Benchmarking results for six methods on the simulated VZV datasets. 2 to 6 VZV strains with 30x -210x coverage and uniform or non-uniform abundances. The x-axis label (Number of strains, div or uni) indicates the number of strains contained in the dataset and whether their abundances are non-uniform (div) or uniform (uni). For example, the x-axis label (2, d10) means a dataset containing 2 strains, with abundance ratios increasing proportionally to the number of strains (e.g., a ratio of 1:2 for 2 strains), where the smallest strain coverage was set to 10x; The x-axis label (2, uni) indicates a dataset containing 2 strains, with uniform abundance, where each strain coverage was set to 30x.

### Experiments on real norovirus data

Noroviruses belong to the genus *Norovirus*, a large and diverse genus in the positive-strand RNA virus family *Caliciviridae*. As a human enteric pathogen, they are highly contagious and cause substantial morbidity across both health care and community settings [34]. We test devider, HaploDMF, iGDA, RVHaplo, Strainline and virCHap on a real norovirus dataset, which consists of seven strains. The true haplotype sequences and the corresponding ONT reads [35] were also used in HaploDMF. Because the sizes of the seven read datasets differ significantly (518–196 525 reads), we followed a similar sampling strategy as used in HaploDMF. We randomly sampled reads from each dataset while maintaining the smallest dataset at 1% abundance, and only reads with a length between 800 bp and 7000bp were retained, as the genome size is about 7.5kb. The pairwise divergence among these haplotypes ranges from 1.19% to 3.62% (**Supplementary Fig. 1d**). The total sequencing depth is approximately 7800x. The sequence with NCBI accession ID MW661279.1 (7551bp) was used as the reference. As shown in **Table 1**, virChap’s recall (82.67%) was 5.21% higher than devider, the second-best, and the read clustering accuracy (96.33%) exceeded devider, the second-best, by 2.56%. Although virCHap’s precision (98.33%) was 0.96% lower than the top-performing tool iGDA, virCHap’s recall was 5.55% higher than iGDA. The number of haplotypes output by virCHap exactly equaled the true number of strains, with an NGA50 of 7532bp bp, which may indicate the output has better contiguity. virCHap also had the second-best EMD. However, virCHap required more computational time and memory in the real norovirus dataset, indicating it needed further optimization in algorithmic efficiency.

**Table 1.**
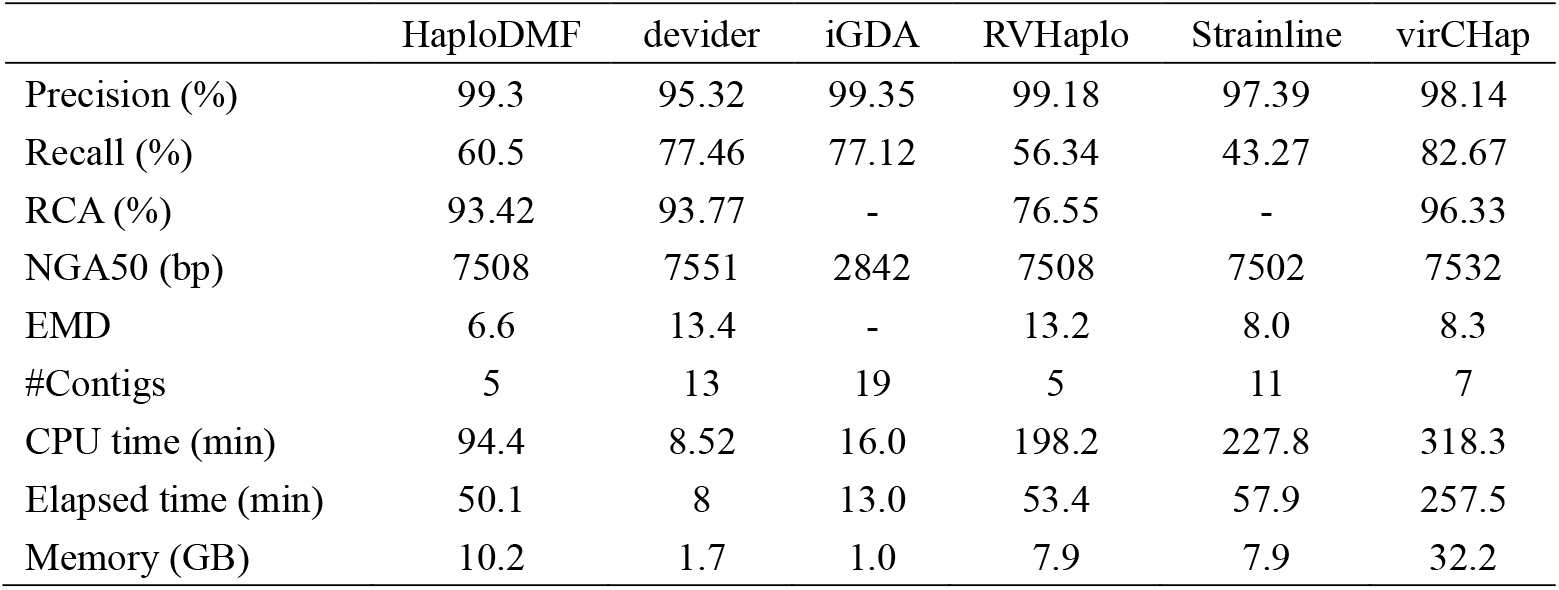
Comparison of devider, HaploDMF, iGDA, RVHaplo, Strainline and virCHap on a real norovirus dataset with 7 strains. RCA: read clustering accuracy; NGA50 is labeled with “-” if the uniquely aligned blocks cover less than half of the reference length; EMD: earth mover’s distance; #Contigs: the number of predicted haplotypes.

|  | HaploDMF | devider | iGDA | RVHaplo | Strainline | virCHap |
| --- | --- | --- | --- | --- | --- | --- |
| Precision (%) | 99.3 | 95.32 | 99.35 | 99.18 | 97.39 | 98.14 |
| Recall (%) | 60.5 | 77.46 | 77.12 | 56.34 | 43.27 | 82.67 |
| RCA (%) | 93.42 | 93.77 | - | 76.55 | - | 96.33 |
| NGA50 (bp) | 7508 | 7551 | 2842 | 7508 | 7502 | 7532 |
| EMD | 6.6 | 13.4 | - | 13.2 | 8.0 | 8.3 |
| #Contigs | 5 | 13 | 19 | 5 | 11 | 7 |
| CPU time (min) | 94.4 | 8.52 | 16.0 | 198.2 | 227.8 | 318.3 |
| Elapsed time (min) | 50.1 | 8 | 13.0 | 53.4 | 57.9 | 257.5 |
| Memory (GB) | 10.2 | 1.7 | 1.0 | 7.9 | 7.9 | 32.2 |

### Experiments on real Potato virus Y (PVY) data

Potato virus Y (PVY), a type member of the genus *Potyvirus* in the family *Potyviridae*, has a single-stranded, positive-sense RNA genome of approximately 9.7 kb. PVY has a wide host range that includes many genera in the family *Solanaceae*, including important crop plants such as potato, pepper, and tobacco [36]. We test various methods on a real potato virus Y (PVY) dataset comprising five haplotype strains. The true haplotype sequences [37] were also used in Strainline. The corresponding real ONT read datasets are available under the study accession number PRJNA612026 [37]. The pairwise divergence among these haplotypes ranges from 3.6% to 21.6% (**Supplementary Fig. 1e**). The abundance ratio of the mixed read datasets is about 8.4:16.8:22.4:24:28.4 and the total sequencing depth is approximately 4100x. The reference sequence was accessed from GenBank under accession number GCA_000862905.1 (9704bp). As shown in **Table 2**, virCHap had a recall (97.47%) that was 8.56% higher than the second-best method, HaploDMF, and the read clustering accuracy (92.45%) that exceeded the second-best method, devider, by 32.05%. The precision of virCHap (92.92%) was 6.70% lower than that of the top-performing method, iGDA, its recall (97.47%) was 20.33% higher than iGDA. virCHap produced the longest NGA50 of 9205bp, and ranked second in terms of EMD (111.4). While the runtime and memory usage of virCHap were higher than those of devider and iGDA, it remained more efficient than those of HaploDMF and Strainline.

**Table 2.** Comparison of devider, HaploDMF, iGDA, RVHaplo, Strainline and virCHap on a real PVY dataset with 5 strains. RCA: read clustering accuracy; NGA50 is labeled with “-” if the uniquely aligned blocks cover less than half of the reference length; EMD: earth mover’s distance; #Contigs: the number of predicted haplotypes.

|  | HaploDMF | devider | iGDA | RVHaplo | Strainline | virCHap |
| --- | --- | --- | --- | --- | --- | --- |
| Precision (%) | 94.37 | 90.38 | 99.62 | 97.46 | 98.01 | 92.92 |
| Recall (%) | 88.91 | 67 | 77.14 | 46.14 | 29.13 | 97.47 |
| RCA (%) | 57.68 | 60.4 | - | 50.42 | - | 92.45 |
| NGA50 (bp) | 9106 | 4230 | 2369 | 5612 | - | 7360 |
| EMD | 149.8 | 127.0 | - | 247.8 | 64.5 | 111.4 |
| #Contigs | 8 | 7 | 15 | 3 | 3 | 13 |
| CPU time (h) | 4.06 | 0.15 | 0.15 | 1.6 | 4.4 | 1.49 |
| Elapsed time (h) | 3.19 | 0.13 | 0.1 | 0.65 | 1.13 | 0.68 |
| Memory (GB) | 12.8 | 3.7 | 2.1 | 4.9 | 21.1 | 10.4 |

### Experiments on real SARS-CoV-2 data

We evaluated devider, HaploDMF, iGDA, RVHaplo, Strainline and virCHap on a real SARS-CoV-2 dataset, which consists of the following six strains: Beta, Eta, Lambda, Omicron, Kappa, and Epsilon. The true haplotype sequences [38] were accessed from the GISAID (https://www.gisaid.org/) database, and the pairwise divergence among these haplotypes ranges from 0.18% to 0.31% (**Supplementary Fig. 1f**). The total sequencing depth is approximately 2600x. The sequence labeled “Wuhan-Hu-1” (NC_045512.2; 29,903bp) was used as the reference.

As shown in **Fig. 5a**, we ran all methods on the full-length SARS-CoV-2 genome. We did not show Strainline as it did not produce results. iGDA and RVhaplo had the highest precision, while virCHap’s precision (the third-best) was 4.21% lower than the highest (iGDA), but virCHap’s recall (the best) was 39.86% higher than iGDA, and 4.36% higher than devider, the second-best one. virCHap also had the highest read clustering accuracy, exceeding the second-highest (RVHaplo) by 20.32%, and virCHap had the lowest EMD. HaploDMF had better contiguity than other methods, but it recovered only four strains **(Supplementary Fig. 2a)**. In terms of the number of predicted contigs, virCHap’s results appeared relatively fragmented **(Supplementary Fig. 2a)**. We aligned these six truth haplotype sequences to the reference and counted the variant sites. We found an average of one variant site per 183 bp, with an uneven distribution of variants **(Supplementary Fig. 2b)**. The average length of the mixed six raw sequencing reads **(Supplementary Fig. 2c)** was approximately 782bp, indicating relatively few variant sites per read. If there were insufficient reliable discriminatory variant sites between reads, we opted not to merge them, which may lead to the fragmentation of our results.

**Fig. 5.**
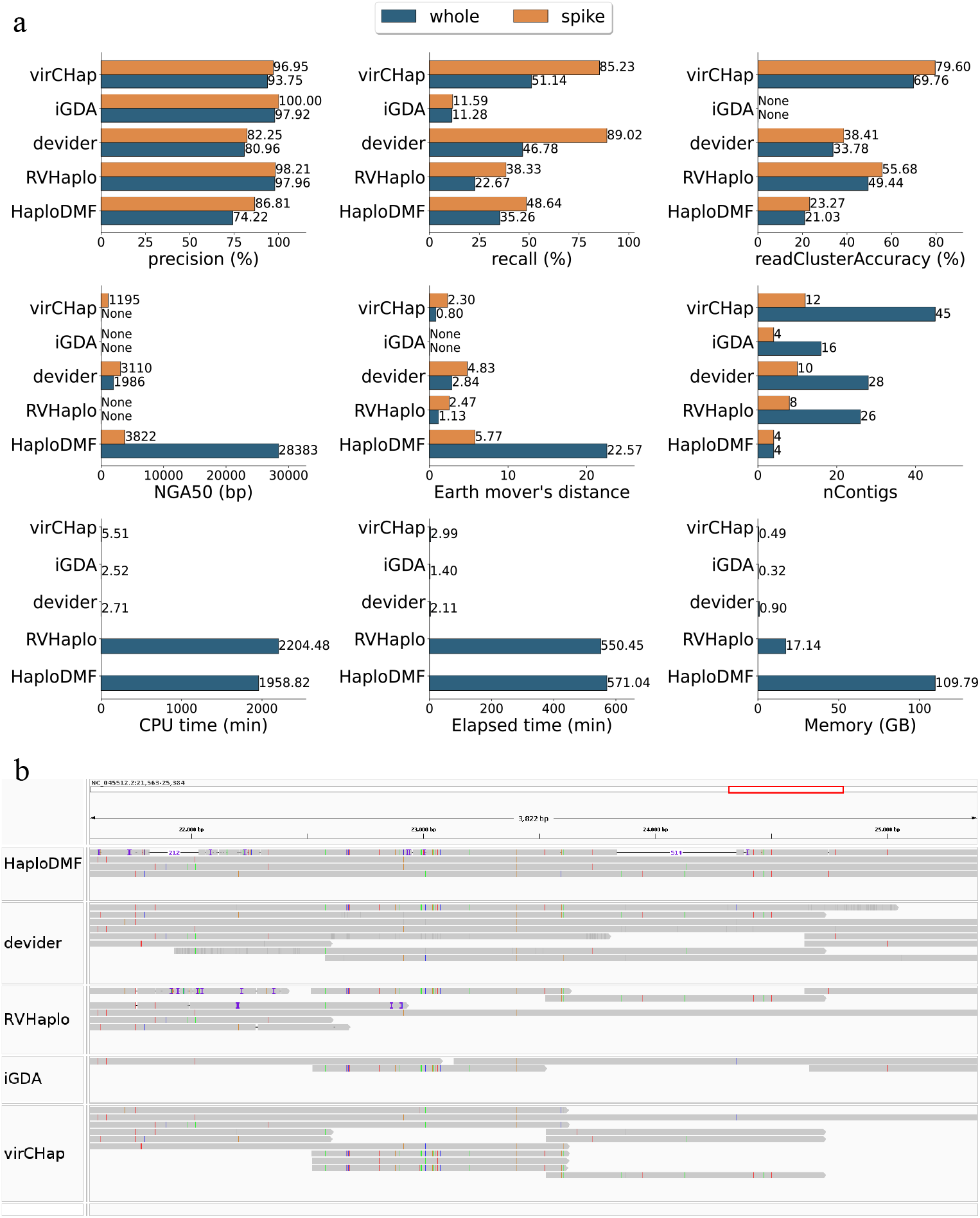
Comparison of devider, HaploDMF, iGDA, RVHaplo and virCHap on a real SARS-CoV-2 dataset with 6 strains. **a**. Various statistics for real SARS-CoV-2 dataset. “whole” means the performance of each method across the entire SARS-CoV-2 genome; “spike” means the performance of each method within the spike gene region of SARS-CoV-2. **b**. IGV [42] screenshot showing predicted results for HaploDMF, devider, RVHaplo, iGDA, and virCHap within the spike gene region of SARS-CoV-2.

As shown in **Supplementary Fig. 2a** and **Supplementary Fig. 2b**, the region between 21 kb and 29 kb had a higher density of variant sites. In this region, there exists the spike gene (NC_045512.2: 21563-25384; about 3.8 kb) [39] of SARS-CoV-2. Nucleotide variants can occur in several genes; notably, major mutations have been observed within the spike gene [40], impacting viral infectivity and host immune response. Furthermore, the spike gene serves as the primary region for classifying variants of concern, which are critical for understanding pandemic origins, transmission dynamics, and disease severity [40] [41]. We counted the variant sites within the spike gene and found that these sites account for approximately one-third of all variant sites (**Supplementary Fig. 2b**), while the spike gene comprises only 12.8% of the entire reference genome length, indicating a higher mutation density in this region. Therefore, we also evaluated the performance of each method specifically within the spike gene (**Fig. 5a**). iGDA had the highest precision, but its recall was the lowest. virCHap had a 14.7% higher precision than devider, although devider’s recall (the best) was 3.79% higher than virCHap (the second-best). Besides, virCHap had the highest read clustering accuracy and the lowest EMD. RVHaplo and HaploDMF require substantially more time and memory than the other three methods. **Fig. 5b** illustrates the predicted haplotypes output by the five methods within the spike gene.

### Comparisons of runtime and memory usage

All benchmarking analyses were performed with 4 threads on a 2.40GHz Intel(R) Xeon(R) Platinum server. To eliminate the effects of varying default input for methods on runtime comparisons, the runtime of raw read alignments was incorporated into the runtimes of HaploDMF, iGDA, and RVHaplo. Similarly, the runtimes of both raw read alignments and LoFreq variant callings were included in the runtimes of divider and virCHap. The CPU time, elapsed time and peak memory usage evaluations for devider, HaploDMF, iGDA, RVHaplo, Strainline and virCHap are available in **Supplementary Fig. 3a∼c**. devider and iGDA were faster than other tools on nearly all benchmark datasets with mean CPU runtimes of 0.25∼7.2 minutes for devider and 1.01∼9.5 minutes for iGDA, and mean elapsed times of 0.2∼6.8 minutes for devider and 0.6∼6.6 minutes for iGDA. They also required less memory, with average usage of 0.1∼2.0 GB for devider and 0.2∼2.5 GB for iGDA.

For the simulated HCV datasets, on average, virCHap took about 83.0 minutes of CPU time and 39.9 minutes of elapsed time with a peak memory of about 11.4GB, while RVhaplo took about 59.1 minutes of CPU time and 15.8 minutes of elapsed time with a peak memory of about 5.4GB, Strainline took about 80.1 minutes of CPU time and 33.4 minutes of elapsed time with a peak memory of about 11.4GB and HaploDMF took about 742.8 minutes of CPU time and 216.0 minutes of elapsed time with a peak memory of about 10.5GB. For the simulated HIV datasets, virCHap was the third fastest after devider and iGDA, taking on average 3.7 minutes of CPU time, 1.6 minutes of elapsed time, 1.9GB of peak memory versus (9.1 minutes, 1.8 minutes, 1.04GB) for RVHaplo, (17.9 minutes, 6.8 minutes, 5.32GB) for HaploDMF and (334.3 minutes, 141.4 minutes, 6.0GB) for Strainline. For the simulated VZV datasets, virCHap had comparable speed to iGDA (1.0 minutes CPU, 0.4 minutes elapsed), with average 1.0 minutes of CPU time, 0.4 minutes of elapsed time, 0.4GB of peak memory versus (16.1 minutes, 3.8 minutes, 2GB) for RVHaplo, (17.7 minutes, 4.7 minutes, 5.3GB) for HaploDMF and (1141.8 minutes, 497.6 minutes, 133.4GB) for Strainline. These statistics implied that virCHap is well applicable in practice.

## Discussion

In this work, we present a method called virCHap for generating viral strain haplotypes from long-read sequencing data. virCHap built an overlap graph from the reduced reads composed of variants and then resolved haplotypes through two-stage clustering, without the need for explicit strain number information. We first clustered highly similar reads to identify reliable compact structures within local and small-scale regions. These clusters were then converted into consensus sequences, which are longer and capable of spanning greater distances, thereby connecting more distant regions. Merging clusters could extend them horizontally to reconstruct longer haplotypes. In genomic regions with sparse variant distribution and excessively short reduced reads support, such as in SARS-CoV-2, virCHap consequently output an insufficient number of supported haplotypes in these regions. virCHap reintroduced the shorter reduced reads, which were initially excluded during the clustering stage, into the subsequent merging step. This strategy compensated for under-supported regions and improved the final results.

By benchmarking virCHap against other methods on both simulated and real datasets, we were able to demonstrate that our method can reconstruct viral haplotypes with high recall and accuracy (>90%) across different viruses, varying coverages, different numbers of strains, distinct genome sizes, and varying abundances, while also achieving the highest read clustering accuracy. In low-coverage scenarios, such as in the HCV experiment where the least abundant strain had only 3x coverage and the total coverage across nine strains was 366x, virCHap still achieved satisfactory results with >98% recall and >98% accuracy, and as coverage increased, both accuracy and recall exceeded 99%. In the challenging HIV datasets with a high strain count and low inter-strain diversity, virCHap maintained high accuracy and good recall. Even on the difficult SARS-CoV-2 data, virCHap was able to recover relatively accurate and complete haplotypes compared to other methods within a short runtime, though with lower haplotype contiguity, which is likely due to the sparse and uneven distribution of variants among SARS-CoV-2 strains. In VZV experiments, we observed that iGDA, devider, and virCHap performed better on uniform abundance datasets, while HaploDMF performed better on non-uniform datasets.

The main limitation of virCHap is that it is a reference-based method, which is highly dependent on reference genomes, the quality of alignment and the subsequent variant calling. Reference-based methods can lead to reference biases, and the reference genomes were not always available for certain viruses. De novo assembly tools can generate novel genome sequences. Reference-based haplotype phasing can be used to complement de novo assembly. Integrating de novo methods with reference-based methods may better leverage the complementary strengths of both and support a more comprehensive view of genetic diversity. However, we still recommend users to use well-assembled de novo contigs or high-quality reference genomes from databases to generate more comprehensive and accurate results.

## Methods

Here, we describe the virCHap algorithm. It takes as input a reference genome, a long-reads BAM file aligned to the reference and a VCF file containing genetic variations called from the long-read BAM file. virCHap utilizes a process (**Fig. 1**): building an overlap graph, clustering reads, and iteratively merging clusters, and it then outputs the phased SNPs, the clustered reads, the base-level sequences for each strain haplotype and their abundance estimates, without prior knowledge of strain haplotype numbers.

### Generating reduced reads and building overlap graphs

Let *n* denote the number of heterozygous variant positions in the reference genome. Each read *r* is represented by a sequence *r*_0_,…, *r*_*n*−1_ over the alphabet ∑ = {*A, C, G,T*, −}, where *r*_*i*_ is the allele base for the *i*th variant position and “−” indicates an uncovered variant, a deletion, or a base that is neither the reference base nor the alternate base listed in the VCF. We also call such reads reduced reads. For two reduced reads *r* and *s*, let *O*(*r, s*) denote the set of overlapping variant positions, and *M*(*r, s*) denote the set of positions within *O*(*r, s*) where they share the same allele bases. The similarity between *r* and *s* is defined as

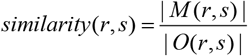

virCHap calculates pairwise similarity scores for all reduced reads and builds an overlap graph *G* = (*V, E*), where each node *v* ∈*V* corresponds to a reduced read. An undirected edge (*r, s*) ∈ *E* is created if the following two conditions are met:

i. | *O* (*r, s*) |≥ *minOverlapLen* (default 25); or 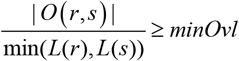 (default 10%) and | *O* (*r, s*) |≥ *minOvlLen* (default 10), where *L*(*r*) is the length of reduced read *r*.
ii. *similarity*(*r, s*) ≥ *minSim* (default 0.95).

The weight of each edge is the similarity between the two reads. Here, we only keep the edges representing highly similar read pairs.

### Clustering and merging clusters

#### Initial clustering via label propagation

First, the overlap graph is partitioned using the label propagation algorithm, a community detection method, which groups reads likely originating from the same haplotype into initial clusters (**Fig. 1B**).

#### Quantifying cluster consensus and merge impact

For a cluster *R*, the proportion of allele base *b* ∈{*A, C, G,T*} at position *i* in *R* is defined as

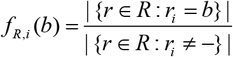

where the denominator is the number of reads in cluster *R* covering position *i*. Let *m*_*R,i*_ denote the major base at position *i* in *R*. For a cluster *R*, let *C*_*R*_ be its consensus sequence, determined by taking the major base at each variant position supported by the reads in *R*. These consensus sequences can be considered an extended-length version of the reduced reads, and can be used to calculate pairwise similarity between the clusters.

Consider merging clusters *R* and *S* into a new cluster *Q* = *R* ∪ *S*, let *N*_*Q,i*_ =|{*r* ∈ *Q* : *r*_*i*_ ≠ −}| be the number of reads covering position *i* in the merged cluster *Q*. We define the merge impact on the original cluster *R* to assess the effect of this union. For each position *i* ∈ *O* (*C*_*R*_, *C*_*Q*_) and allele *b* ∈{*A, C, G,T*}, the change in support, Δ_*R*→*Q*_ (*i, b*), is calculated only under two specific conditions:

i. If *b* = *m*_*R,I*_ and *f*_*Q,i*_ (*b*) < *f*_*R,i*_ (*b*) :

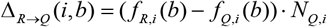
ii. If *b* ≠ *m*_*R,i*_ and *f*_*Q,i*_ (*b*) > *f*_*R,i*_ (*b*) :

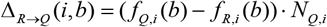

Otherwise, Δ_*R*→*Q*_ (*i, b*) = 0 .

The merging impact on *R* is defined as

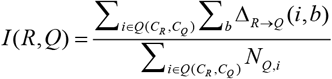

This metric quantifies the average negative impact of merging on *R*. The impact *I*(*S, Q*) on cluster *S* is defined analogously.

#### Hierarchical merging of clusters

Clusters are merged using a bottom-up strategy similar to hierarchical clustering. Consensus sequence pairs (representing the cluster pairs) are sorted in descending order of similarity. In each iteration, the most similar pair (*R, S*) is evaluated. If both *I*(*R, Q*) and *I*(*S, Q*) are below a threshold (default 0.01), the clusters are merged (**Fig. 1C** case 1). The new cluster *Q* replaces *R* and *S*, similarities between *Q* and all adjacent clusters are recomputed, and the pair ordering is updated. If either *I*(*R, Q*) or *I*(*S, Q*) exceeds the threshold, the merge is rejected (**Fig. 1C** case 2) and this pair is removed from the ordering set. The process continues until no remaining pair can be merged.

For genomic regions with sparse and unevenly distributed variants coupled with short aligned read support, such as in SARS-CoV-2, the reduced reads are often short and contain few variant sites. In such cases, virCHap detects regions with insufficient cluster support from the label propagation clustering stage. It then reduces the thresholds *minOvl* and *minOvlLen* to rescue previously excluded short reduced reads. Each of these recovered reads is treated as a separate cluster and they are subsequently merged with the primary clusters produced in the label propagation clustering step. This mechanism is triggered specifically when sparse variant sites and short read length lead to minimal and unreliable overlaps between reduced reads.

#### Output Generation

Upon completing merging, each remaining consensus corresponds to a haplotype cluster, containing several reads. The abundance of a haplotype is estimated as the fraction of assigned reads. Haplotypes with low abundance and low sequencing depth are filtered out, and the consensus sequences of the remaining haplotypes are the final phased SNPs. Finally, base-level consensus sequences are generated for each haplotype using the assigned reads and the corresponding BAM file.

## Data availability

All scripts needed to handle the input data and to reproduce the analyses can be found at https://github.com/gaoyun-25/virCHap-test. The haplotype sequences for the HCV strains are available with the accession ID [30]: EU255965, EU255973, EU255980, EU255981, EU255982, EU155339, EU155344, EU234065, EU255983, and EU255989. The haplotype sequences for the HIV strains are available with the accession ID [32]: OR484006, OR483988, OR484012, OR483999, OR483976, OR483982, OR483994, OR483989, OR483985, OR484004, OR484024, OR466100, OR466093, OR483977, OR466094, OR466113, OR483980, OR483979, OR483970, OR454050, and OR483991. The haplotype sequences for the VZV strains are available with the NCBI accession ID: MH709317, DQ479960, DQ479955, AY548171, KF811485, DQ479954, and NC_001348. The haplotype sequences for the norovirus strains are available with the accession ID [35]: MW661283, MW661278, MW661264, MW661261, MW661260, MW661259, MW661258 and MW661279. The corresponding Oxford Nanopore data for the norovirus strains is available with the SRA accessions [35]: SRR13951160, SRR13951165, SRR13951181, SRR13951184, SRR13951185, SRR13951186, and SRR15525305. The haplotype sequences for the PVY strains are available with the accession ID [37]: MT264732, MT264731, MT264735, MT264734, MT264733, and NC_001616. The corresponding Oxford Nanopore data for the PVY strains is available with the SRA accessions [37]: SRR11431596, SRR11431597, SRR11431615, SRR11431616, and SRR11431617. The haplotype sequences for the SARS-CoV-2 are available at GISAID (https://www.gisaid.org/) with accession ID [38]: EPI_ISL_1615591, EPI_ISL_1660405, EPI_ISL_1494722, EPI_ISL_6814922, EPI_ISL_1660480, and, EPI_ISL_717710. The corresponding Oxford Nanopore data for the SARS-CoV-2 strains is available with the SRA accessions [38]: SRR18680774, SRR18680775, SRR18680776, SRR18680779, SRR18680780, and SRR18680782.

## Code availability

The source code for the latest version of the virCHap package is available at https://github.com/gaoyun-25/virCHap.

## Funding

This work was supported by the National Key R&D Program of China with code 2020YFA0712400; the Fundamental Research Funds for the Central Universities; the National Natural Science Foundation of China with codes 12471461 and 11931008. The funders had no role in the study design, data collection and analysis, decision to publish, or preparation of the manuscript.

## Authors’ contributions

Y.G. and T.Y. conceived and designed the experiments. Y.G. performed the experiments. Y.G. contributed reagents/materials/analysis tools. Y.G. wrote the paper. Y.G. and T.Y. designed the software used in the analysis. G.L., B.L., and T.Y. oversaw the project.

## Competing interest statement

The authors declare that they have no competing interests.

## Notes

### Competing Interest Statement

The authors have declared no competing interest.

## Reference

1. Duarte, E.A., et al., RNA virus quasispecies: significance for viral disease and epidemiology. Infect Agents Dis, 1994. 3(4): p. 201–14.

2. Domingo, E., J. Sheldon, and C. Perales, Viral Quasispecies Evolution. Microbiology and Molecular Biology Reviews, 2012. 76(2): p. 159–216.

3. Lauring, A.S. and R. Andino, Quasispecies Theory and the Behavior of RNA Viruses. PLOS Pathogens, 2010. 6(7): p. e1001005.

4. Grubaugh, N.D., et al., Tracking virus outbreaks in the twenty-first century. Nature Microbiology, 2019. 4(1): p. 10–19.

5. Popa, A., et al., Genomic epidemiology of superspreading events in Austria reveals mutational dynamics and transmission properties of SARS-CoV-2. Science Translational Medicine, 2020. 12(573): p. eabe2555.

6. Lu, J., et al., Genomic Epidemiology of SARS-CoV-2 in Guangdong Province, China. Cell, 2020. 181(5): p. 997-1003.e9.

7. Koren, S., et al., Canu: scalable and accurate long-read assembly via adaptive k-mer weighting and repeat separation. Genome Res, 2017. 27(5): p. 722–736.

8. Kolmogorov, M., et al., Assembly of long, error-prone reads using repeat graphs. Nature Biotechnology, 2019. 37(5): p. 540–546.

9. Shafin, K., et al., Nanopore sequencing and the Shasta toolkit enable efficient de novo assembly of eleven human genomes. Nature Biotechnology, 2020. 38(9): p. 1044–1053.

10. Cheng, H., et al., Haplotype-resolved de novo assembly using phased assembly graphs with hifiasm. Nature Methods, 2021. 18(2): p. 170–175.

11. Nurk, S., et al., HiCanu: accurate assembly of segmental duplications, satellites, and allelic variants from high-fidelity long reads. Genome Res, 2020. 30(9): p. 1291–1305.

12. Feng, X., et al., Metagenome assembly of high-fidelity long reads with hifiasm-meta. Nature Methods, 2022. 19(6): p. 671–674.

13. Benoit, G., et al., High-quality metagenome assembly from long accurate reads with metaMDBG. Nature Biotechnology, 2024. 42(9): p. 1378–1383.

14. Shaw, J., et al., Long-read reconstruction of many diverse haplotypes with devider. Genome Res, 2025. 35(12): p. 2637–2649.

15. Eliseev, A., et al., Evaluation of haplotype callers for next-generation sequencing of viruses. Infection, Genetics and Evolution, 2020. 82: p. 104277.

16. Mantere, T., S. Kersten, and A. Hoischen, Long-Read Sequencing Emerging in Medical Genetics. Frontiers in Genetics, 2019. Volume 10 -2019.

17. Zagordi, O., et al., ShoRAH: estimating the genetic diversity of a mixed sample from next-generation sequencing data. BMC Bioinformatics, 2011. 12(1): p. 119.

18. Ahn, S. and H. Vikalo, aBayesQR: A Bayesian Method for Reconstruction of Viral Populations Characterized by Low Diversity. Journal of Computational Biology, 2018. 25(7): p. 637–648.

19. Knyazev, S., et al., Accurate assembly of minority viral haplotypes from next-generation sequencing through efficient noise reduction. Nucleic Acids Research, 2021. 49(17): p. e102–e102.

20. Feng, Z., et al., Detecting and phasing minor single-nucleotide variants from long-read sequencing data. Nature Communications, 2021. 12(1): p. 3032.

21. Cai, D. and Y. Sun, Reconstructing viral haplotypes using long reads. Bioinformatics, 2022. 38(8): p. 2127–2134.

22. Cai, D., J. Shang, and Y. Sun, HaploDMF: viral haplotype reconstruction from long reads via deep matrix factorization. Bioinformatics, 2022. 38(24): p. 5360–5367.

23. Luo, X., X. Kang, and A. Schönhuth, Strainline: full-length de novo viral haplotype reconstruction from noisy long reads. Genome Biology, 2022. 23(1): p. 29.

24. Raghavan, U.N., R. Albert, and S. Kumara, Near linear time algorithm to detect community structures in large-scale networks. Physical Review E, 2007. 76(3): p. 036106.

25. Wick, R., Badread: simulation of error-prone long reads. Journal of Open Source Software, 2019. 4: p. 1316.

26. Li, H., Minimap2: pairwise alignment for nucleotide sequences. Bioinformatics, 2018. 34(18): p. 3094–3100.

27. Wilm, A., et al., LoFreq: a sequence-quality aware, ultra-sensitive variant caller for uncovering cell-population heterogeneity from high-throughput sequencing datasets. Nucleic Acids Research, 2012. 40(22): p. 11189–11201.

28. Mikheenko, A., V. Saveliev, and A. Gurevich, MetaQUAST: evaluation of metagenome assemblies. Bioinformatics, 2016. 32(7): p. 1088–1090.

29. Bartenschlager, R., V. Lohmann, and F. Penin, The molecular and structural basis of advanced antiviral therapy for hepatitis C virus infection. Nature Reviews Microbiology, 2013. 11(7): p. 482–496.

30. Kuntzen, T., et al., Naturally occurring dominant resistance mutations to hepatitis C virus protease and polymerase inhibitors in treatment-naïve patients. Hepatology, 2008. 48(6).

31. Jain, C., et al., High throughput ANI analysis of 90K prokaryotic genomes reveals clear species boundaries. Nature Communications, 2018. 9(1): p. 5114.

32. Kinloch Natalie, N., et al., HIV reservoirs are dominated by genetically younger and clonally enriched proviruses. mBio, 2023. 14(6): p. e02417–23.

33. Liu, J., et al., Genotyping of Clinical Varicella-Zoster Virus Isolates Collected in China. Journal of Clinical Microbiology, 2009. 47(5): p. 1418–1423.

34. Robilotti, E., S. Deresinski, and A. Pinsky Benjamin, Norovirus. Clinical Microbiology Reviews, 2015. 28(1): p. 134–164.

35. Flint, A., et al., Genomic analysis of human noroviruses using combined Illumina–Nanopore data. Virus Evolution, 2021. 7(2): p. veab079.

36. Karasev, A.V. and S.M. Gray, Continuous and emerging challenges of Potato virus Y in potato. Annu Rev Phytopathol, 2013. 51: p. 571–86.

37. Della Bartola, M., S. Byrne, and E. Mullins Characterization of Potato Virus Y Isolates and Assessment of Nanopore Sequencing to Detect and Genotype Potato Viruses. Viruses, 2020. 12, 478 DOI: 10.3390/v12040478.

38. Aggarwal, A., et al., Platform for isolation and characterization of SARS-CoV-2 variants enables rapid characterization of Omicron in Australia. Nature Microbiology, 2022. 7(6): p. 896–908.

39. Liao, Y.-C., et al. High-Integrity Sequencing of Spike Gene for SARS-CoV-2 Variant Determination. International Journal of Molecular Sciences, 2022. 23, 3257 DOI: 10.3390/ijms23063257.

40. Nimsamer, P., et al., “Nano COVID-19”: Nanopore sequencing of spike gene to identify SARS-CoV-2 variants of concern. Exp Biol Med (Maywood), 2023. 248(20): p. 1841–1849.

41. Harvey, W.T., et al., SARS-CoV-2 variants, spike mutations and immune escape. Nature Reviews Microbiology, 2021. 19(7): p. 409–424.

42. Robinson, J.T., et al., Integrative genomics viewer. Nature Biotechnology, 2011. 29(1): p. 24–26.

